# CA19-9 promotes liver metastasis of pancreatic cancer through E-selectin mediated extravasation

**DOI:** 10.64898/2026.04.08.717301

**Authors:** Satoshi Ogawa, Hyemin Song, Jasper Hsu, Vasiliki Pantazopoulou, Victoria Osorio-Vasquez, Casie S. Kubota, Jacob R. Tremblay, Chelsea R. Bottomley, Kathryn Lande, Jonathan Zhu, Kristina L. Peck, Yongjia Wang, Kassidy Curtis, Sophie Keightley, Ryan Tomita, Jingjing Zou, Michael Downes, Ronald M. Evans, Andrew M. Lowy, Herve Tiriac, Dannielle D. Engle

## Abstract

Pancreatic ductal adenocarcinoma (PDAC) frequently metastasizes to the liver, which drives patient mortality. CA19-9 is elevated in most PDAC tumors and is widely used as a clinical biomarker. Elevated serum levels are associated with poor outcomes.

However, whether CA19-9 functionally contributes to metastatic progression has not been fully defined, in part because mice lack endogenous CA19-9 expression. Here, using syngeneic murine PDAC cells engineered to express CA19-9, we investigated its functional role in liver metastasis. In splenic injection models, CA19-9 expression markedly increased liver metastatic burden by promoting both metastatic seeding and subsequent metastatic outgrowth. *In vitro*, CA19-9 enhanced tumor cell adhesion to endothelial cells through interaction with E-selectin. Metastatic seeding of CA19-9-expressing cells was reduced by genetic deletion of E-selectin or antibody neutralization of either CA19-9 or E-selectin *in vivo*. Therapeutic targeting of CA19-9 with a neutralizing antibody markedly reduced liver metastatic burden after metastatic seeding. CA19-9 expression increased AKT signaling in PDAC cells and liver metastases, and CA19-9 levels correlated with AKT activation in human PDAC tissues. These findings show that CA19-9 promotes PDAC liver metastasis through E-selectin-dependent metastatic seeding and AKT-associated metastatic outgrowth, highlighting CA19-9 as a functional mediator of PDAC metastasis and a potential therapeutic target.

## Introduction

Pancreatic ductal adenocarcinoma (PDAC) remains one of the most lethal malignancies with a five-year survival rate of 13% in the United States (1). The prognosis of patients with distant metastasis is particularly poor, with survival rates falling to 3% (1). Unfortunately, most patients will be diagnosed with PDAC after it has already metastasized (2, 3). The most common site of metastasis is the liver (4). The dismal outcomes for patients with metastatic PDAC highlight the limited efficacy of standard of care combination chemotherapy once patients reach this stage. Even among patients with initially resectable disease, recurrence is frequent, most commonly in the liver, where it is associated with poor survival outcomes (5). Given the current clinical reality, more effective treatment options for metastatic PDAC are essential.

Metastasis is a multistep process involving tumor cell migration, intravasation, survival in circulation, extravasation, and colonization at distant organs (6). A deeper understanding of these sequential steps and the molecular mechanisms that regulate tumor cell behavior *in vivo* is essential for the development of effective therapeutic strategies. Thus, identifying how specific tumor-associated molecules contribute to distant metastasis is particularly important for improving outcomes in PDAC.

Glycosylation, a common post-translational modification, is often dysregulated in cancer, resulting in increased expression of glycoproteins that can serve as tumor markers and modulators of tumor behavior (7, 8). Aberrant glycosylation has been implicated in promoting tumor progression and metastasis through mechanisms such as modulation of growth factor receptor signaling, alteration of cell-cell and cell-matrix interactions, and facilitation of immune evasion (8, 9). These glycosylation-dependent processes influence the multistep metastatic cascade, shaping how tumor cells establish distant metastasis. In this context, understanding the contributions of glycan-mediated pathways is crucial for elucidating the molecular mechanisms of PDAC metastasis and provides new opportunities for therapeutic interception.

The glycan CA19-9 (sialyl Lewis a: sLe^a^) is the most widely used tumor marker for PDAC. CA19-9 is generated by the sequential addition of sugars to type I precursor chains present on proteins and other molecules (10). The critical final step is the α1,4 linkage of fucose to N-acetylglucosamine, a reaction catalyzed by fucosyltransferase 3 (FUT3), which is the unique enzyme capable of producing CA19-9 (11, 12). Elevated serum CA19-9 is observed in approximately 70-90% of PDAC and is useful for tracking treatment response, but not early detection due to its limited specificity (13, 14).

Importantly, multiple retrospective clinical studies have reported significant associations between elevated preoperative serum CA19-9 levels and poor prognosis or early postoperative recurrence in PDAC patients (13, 15–19). Consistent with these observations, systematic reviews and multiple meta-analyses have confirmed that preoperative serum CA19-9 is not only a biomarker, but also a prognostic factor for early recurrence after pancreatic cancer resection (20). Although whether CA19-9 directly contributes to PDAC progression or merely reflects tumor burden remains unclear, these clinical associations raise the possibility that CA19-9 may have functional relevance in PDAC metastasis.

Previous studies have shown that CA19-9 is a ligand for E-selectin, an endothelial adhesion molecule that is induced during inflammation. *In vitro* assays using CA19-9-positive cells and endothelial cells have demonstrated that CA19-9-modified structures can bind E-selectin (21–23). These studies have suggested that CA19-9 may facilitate metastatic dissemination by promoting interaction with E-selectin in the vasculature.

However, studying its biological role in PDAC metastasis *in vivo* remains elusive because mice do not produce CA19-9 due to the lack of Fut3, which is a pseudogene in rodents (24, 25). Our group has addressed this gap by creating syngeneic murine PDAC cell lines and mouse models expressing CA19-9 through ectopic expression of B3GALT5 and FUT3. These models enable human-like expression of this glycan and have demonstrated that CA19-9 promotes pancreatitis and pancreatic cancer in mice (26). Here, we leveraged these models to investigate the functional role of CA19-9 in liver metastasis of PDAC *in vivo*, revealing potential therapeutic opportunities.

Therapeutic strategies targeting the CA19-9-E-selectin axis have been explored. A humanized monoclonal antibody against CA19-9 (5B1) and pharmacologic inhibition of E-selectin using small-molecule antagonists such as Uproleselan (GMI-1271) have been evaluated in early-phase clinical studies, demonstrating acceptable safety and biological activity in patients with CA19-9-expressing malignancies (27) or hematologic cancers (28). However, despite their clinical feasibility, the therapeutic relevance of targeting CA19-9 or E-selectin in PDAC, particularly in the context of metastatic progression, remains insufficiently defined. Given the frequent occurrence of liver metastasis and early postoperative recurrence in PDAC, elucidating whether and how the CA19-9-E-selectin axis contributes to liver metastasis *in vivo* is important.

## Results

### CA19-9 promotes PDAC liver metastasis

To express CA19-9 in murine pancreatic cancer cells, we transduced FC1199 and FC1245, derived from Kras^LSL-G12D/+^; Trp53^LSL-R172H/+^; Pdx1-Cre (KPC) mice, and KPCY 6419c5 (KPCY), derived from KPC mice harboring a YFP reporter, with human *FUT3* and *B3GALT5* (Supplemental Figure 1A) (26). To determine whether CA19-9 affects the growth and migration of pancreatic cancer cells, we first performed proliferation and scratch assays. CA19-9 expression did not alter the proliferation of FC1199 or KPCY cells, indicating that CA19-9 does not directly influence cell growth in nutrient replete conditions (Supplemental Figure 2A, B). In contrast, CA19-9^pos^ cells exhibited significantly accelerated wound closure in scratch assays (Supplemental Figure 2C-E). These results indicate that CA19-9 enhances migratory capacity without affecting basal proliferation.

To investigate the role of CA19-9 in pancreatic cancer liver metastasis *in vivo*, we employed a hemisplenic injection model, which allows tumor cells to directly enter the portal circulation and efficiently seed the liver, followed by a hemisplenectomy. Using this approach, we injected FC1199 CA19-9^neg^ and CA19-9^pos^ cells. Splenic injection of FC1199 CA19-9^pos^ cells resulted in a markedly higher liver metastatic burden compared to CA19-9^neg^ cells (Figure 1A, B). The liver weight and liver-to-body weight ratio were significantly increased in mice injected with CA19-9^pos^ cells (Figure 1C), indicating higher metastatic burden. Hematoxylin and eosin staining (H&E) revealed comparable tumor morphology between the CA19-9^neg^ and CA19-9^pos^ tumors (Figure 1D).

**Figure 1.**
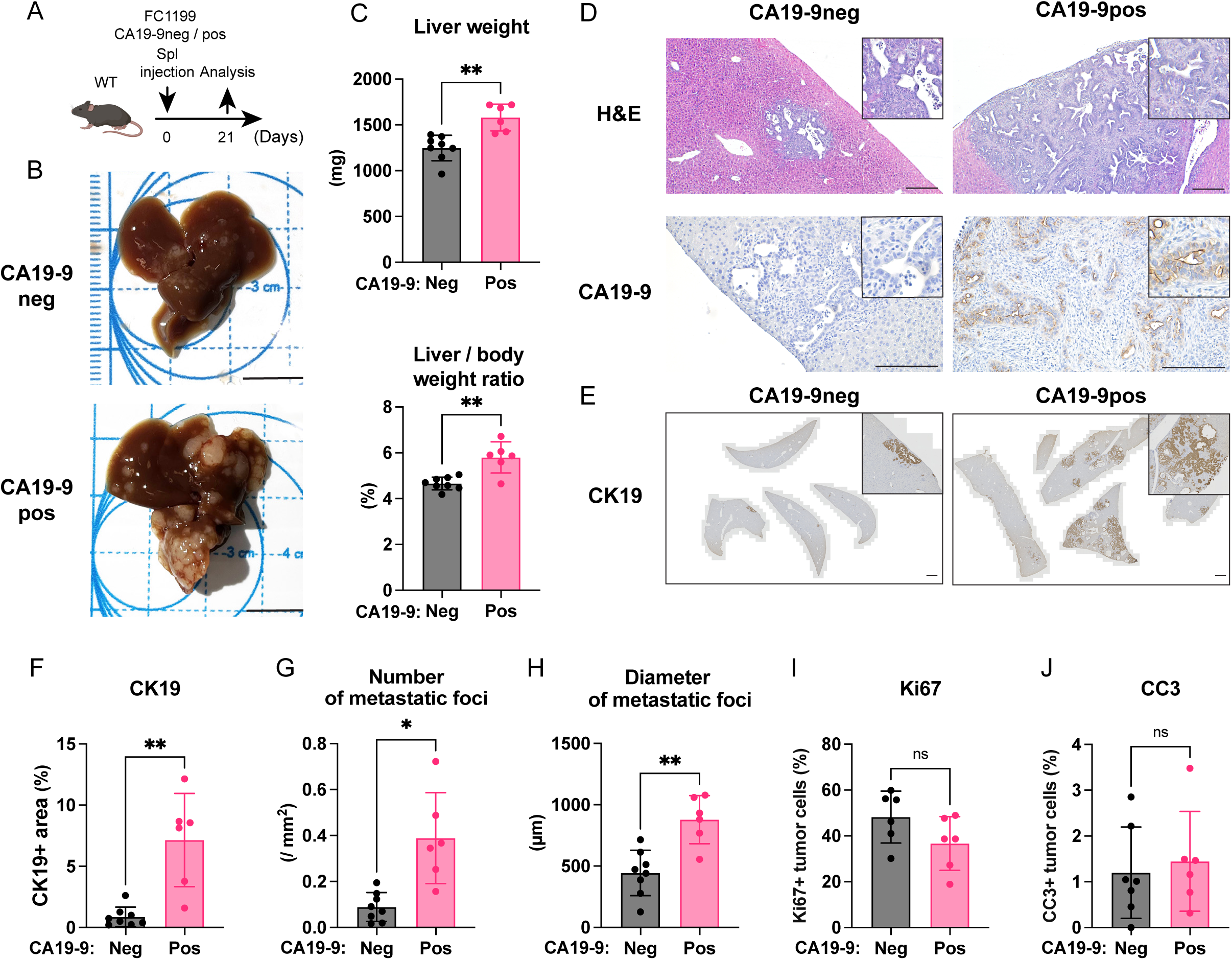
CA19-9-positive cells promote the seeding and outgrowth of liver metastasis. (A) Schema of study design in (B-J). (B) Representative macroscopic images of the livers isolated from mice injected with CA19-9^neg^ (top) or CA19-9^pos^ (bottom) cells. Scale bar = 1 cm. (C) Quantification of liver weight (top) and of liver weight normalized by body weight (bottom). (D) Representative H&E (top row) and CA19-9 IHC (bottom row) images of the liver metastasis in splenic injection models of CA19-9^neg^ (left) and CA19-9^pos^ (right) cells. Scale bar = 200 μm. (E) CK19 IHC of whole liver sections in splenic injection models of CA19-9 negative (left) and positive (right) cells. Scale bar = 1 mm. (F) Quantification of CK19 positive area (%) across the whole liver sections. (G) Number of metastatic foci across the whole liver sections normalized by tissue area (mm²). (H) Mean diameter of metastatic foci measured across whole liver sections. (I, J) Quantification of Ki67 positive (I) and cleaved caspase-3 (CC3) positive (J) tumor cells within liver metastasis area. *Data are presented as mean ± SD. Each dot represents an individual mouse. Mice were injected with 1 × 10^5^ tumor cells per mouse. Sample sizes may vary between analyses because some liver sections lacked detectable metastatic lesions. Statistical significance was determined by unpaired two-tailed t-test with Welch’s correction. *P < 0.05; **P < 0.01; ns, not significant.

Immunohistochemistry (IHC) for CA19-9 clearly showed CA19-9^pos^ tumors exhibited strong membranous and luminal staining patterns compared with CA19-9^neg^ tumors (Figure 1D). IHC for CK19 demonstrated metastatic tumors occupied a significantly larger area across whole liver sections of mice injected with CA19-9^pos^ cells (Figure 1E, F). Both the number and diameter of metastatic foci were significantly higher in the livers of mice injected with CA19-9^pos^ cells (Figure 1G, H). The increase in the number of metastatic foci can reflect enhanced metastatic seeding in the liver while the larger diameter of metastatic foci is often indicative of elevated outgrowth. To assess the proliferative and apoptotic status of metastases, we performed IHC for Ki67 and cleaved caspase-3 (CC3). No significant difference in either proliferation or apoptosis was found between CA19-9^neg^ and CA19-9^pos^ metastatic lesions (Figure 1I, J, Supplemental Figure 3A). Significantly increased liver metastases were also found following splenic injection of CA19-9^pos^ KPCY cells compared with CA19-9^neg^, consistent with the findings in FC1199 cells (Supplemental Figure 3B-F). Similarly, Ki67 and CC3 staining showed no significant differences between the groups (Supplemental Figure 3G, H). Taken together, these data demonstrate that CA19-9 promotes not only the seeding of metastatic foci but also subsequent outgrowth of PDAC liver metastasis.

### CA19-9 enhances adhesion to endothelial cells through E-selectin *in vitro*

Vascular adhesion is a key step in extravasation during the metastatic process. To determine if CA19-9 promotes metastasis by affecting vascular adhesion, we assessed the adhesion of pancreatic cancer cells to Human Umbilical Vein Endothelial Cells (HUVECs) (21–23). HUVECs upregulate E-selectin expression in response to inflammatory stimuli such as TNF-α (21–23). To model this, HUVECs were pretreated with TNF-α to induce E-selectin expression, followed by incubation with tumor cells to quantify adhesion. Calcein-AM stained CA19-9^pos^ PDAC cells (FC1199, FC1245 and KPCY) exhibited a robust increase in adhesion to endothelial cells compared with CA19-9^neg^ cells (Figure 2A, B), suggesting that CA19-9 facilitates initial vascular adhesion, thereby promoting extravasation. Given the known interaction of CA19-9 and E-selectin (21–23), we next tested whether blocking this interaction could suppress adhesion of CA19-9^pos^ FC1199 cells. Treatment with anti-CA19-9 antibody (5B1) significantly reduced the number of adhered CA19-9^pos^ cells (Figure 2C). Similarly, blockade of endothelial E-selectin using anti-E-selectin antibody (BBA16) markedly decreased the adhesion of CA19-9^pos^ cells (Figure 2D). Treatment of human CA19-9^pos^ PDAC cells (hM19a 2D and SW1990) expressing CA19-9 (Supplemental Figure 1B) with anti-CA19-9 or anti-E-selectin antibodies also decreased adhesion to HUVECs (Figure 2E, F and Supplemental Figure 4). Additionally, to assess the effect of endogenous CA19-9, we generated *FUT3*-knockout Capan-2 cells, as FUT3 is required for CA19-9 biosynthesis. Loss of CA19-9 expression was confirmed by immunoblotting (Supplemental Figure 1C). Loss of FUT3 significantly reduced adhesion of Capan-2 cells to HUVECs (Figure 2G). These findings suggest that CA19-9 promotes tumor cell adhesion to endothelial cells through E-selectin *in vitro*, which may facilitate metastatic dissemination *in vivo*.

**Figure 2.**
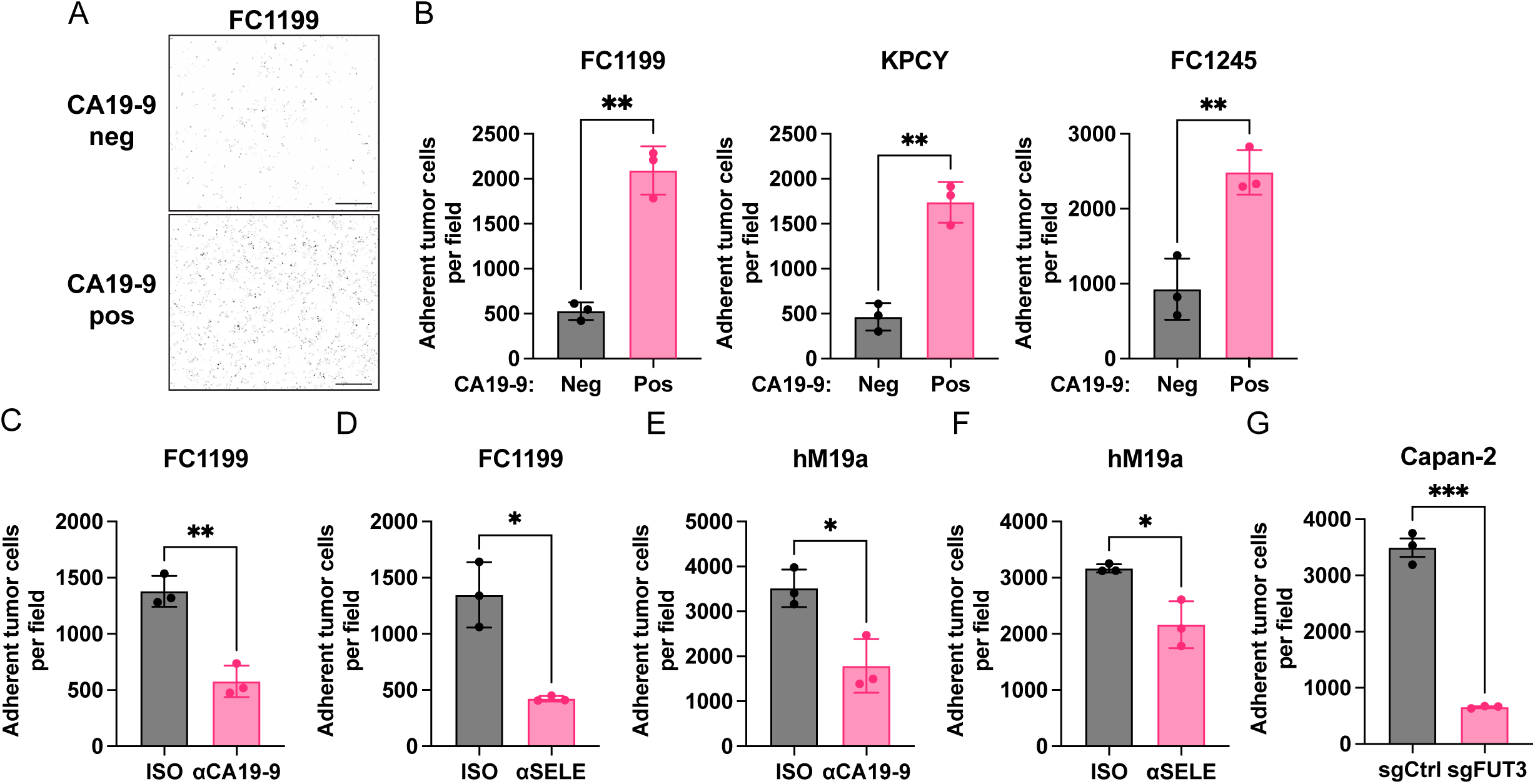
CA19-9 promotes PDAC cell adhesion to endothelial cells through E-selectin binding *in vitro*. (A) Representative images of adhesion assay using Calcein-AM stained CA19-9 negative (top) and positive (bottom) cells adhered to HUVECs. Scale bars = 100 μm. (B) Quantification of adhered CA19-9^neg^ and CA19-9^pos^ FC1199 (left), KPCY (middle) and FC1245 (right) cells per field. (C, D) Quantification of adhered CA19-9^pos^ FC1199 cells treated with anti-CA19-9 antibody (5B1; αCA19-9) (C) and anti-human E-selectin antibody (BBA16; αSELE) (D) compared with isotype control (ISO). (E, F) Quantification of adhered hM19a 2D cells treated with anti-CA19-9 antibody (5B1; αCA19-9) (E) and anti-human E-selectin antibody (BBA16; αSELE) (F) compared with isotype control (ISO). (G) Quantification of adhered Capan-2 cells after FUT3 knockout (sgFUT3) compared with a negative control (sgCtrl). *Data are presented as mean ± SD. Data are representative of at least two independent experiments. Statistical significance was determined by unpaired two-tailed t-test with Welch’s correction. *P < 0.05; **P < 0.01; ***P < 0.001.

### CA19-9 promotes metastatic seeding via E-selectin binding *in vivo*

We next sought to evaluate the functional role of CA19-9 on metastatic seeding of the liver *in vivo*. To determine whether CA19-9 enhances the seeding of liver metastasis, we analyzed the livers of mice one day following splenic injection of CA19-9^neg^ or CA19-9^pos^ KPCY cells (Figure 3A). IHC showed a marked increase in the number of YFP⁺ tumor cells in the liver of mice injected with CA19-9^pos^ compared with CA19-9^neg^ cells (Figure 3B, C), suggesting that CA19-9 promotes initial seeding *in vivo*.

**Figure 3.**
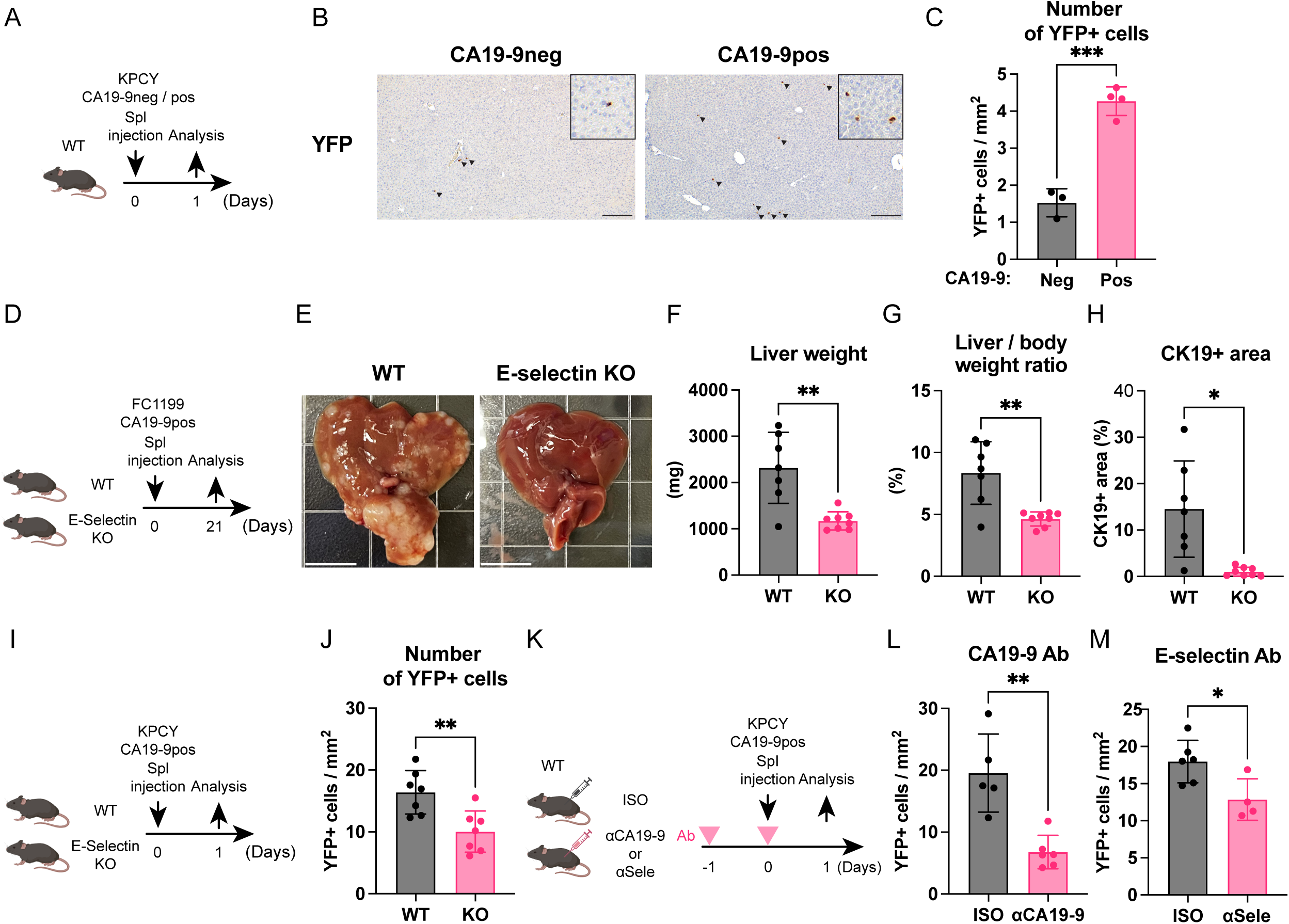
CA19-9 promotes liver metastasis seeding via E-selectin binding *in vivo*. (A) Schema of study design for metastatic seeding analysis in (B, C). (B) Representative YFP IHC of liver sections one day after splenic injection of CA19-9^neg^ or CA19-9^pos^ KPCY cells. Scale bar = 200 μm. (C) Quantification of YFP+ tumor cells in liver sections of WT mice injected with CA19-9^neg^ or CA19-9^pos^ KPCY cells, normalized by tissue area (mm²). (D) Schema of study design in (E-H). (E) Representative macroscopic images of livers isolated from WT (left) and E-selectin KO (right) mice injected with CA19-9^pos^ cells. Scale bar = 1 cm. (F, G) Quantification of liver weight (F) and of liver weight normalized by body weight (G). (H) Quantification of CK19 positive area (%) across the whole liver sections. (I) Schema of study design in (J) (J) Quantification of YFP+ tumor cells in liver sections of WT and E-selectin KO mice injected with CA19-9^pos^ KPCY cells, normalized by tissue area (mm²). (K) Schema of study design in (L, M) (L, M) Quantification of CA19-9^pos^ KPCY cells in liver sections of WT mice injected with CA19-9^pos^ KPCY cells and treated with isotype control or anti-CA19-9 antibody (5B1; αCA19-9) (L), or isotype control or anti-mouse E-selectin antibody (9A9; αSele) (M), normalized by tissue area (mm²). *Data are presented as mean ± SD. Each dot represents an individual mouse. For A-H, mice were injected with 1 × 10^5^ tumor cells per mouse. For I-M, mice were injected with 3 × 10^5^ tumor cells per mouse. Statistical significance was calculated using unpaired two-tailed t-test with Welch’s correction. *P < 0.05; **P < 0.01; ***P < 0.001.

Previous literature suggested that murine E-selectin can bind to CA19-9 (35). To compare the binding capacity of human and murine E-selectin to CA19-9, we generated conditioned media from CA19-9^neg^ and CA19-9^pos^ FC1199 cells, concentrated the supernatants, and performed pull-down assays using recombinant human or murine E-selectin-Fc fusion proteins. CA19-9 immunoblotting revealed comparable binding to both human and murine E-selectin-Fc proteins (Supplemental Figure 5). We sought to determine whether the interaction of CA19-9^pos^ PDAC cells with E-selectin promotes liver metastasis using E-selectin knockout mice (KO; Sele-/-). E-selectin KO mice have no baseline defects in major organs (36). Histological analyses revealed no gross abnormalities in the liver relative to wild-type C57Bl/6J control mice (WT) (Supplemental Figure 6). We analyzed the livers of wild-type or E-selectin KO mice 21 days after splenic injection of CA19-9^pos^ FC1199 cells (Figure 3D). E-selectin KO mice developed substantially fewer liver metastases than wild-type controls (Figure 3E). Both liver weight and liver/body weight ratio were significantly reduced in E-selectin KO mice (Figure 3F, G), and the CK19-positive area was markedly diminished across whole-liver sections (Figure 3H). Because the reduced metastatic burden observed in E-selectin KO mice could arise from either impaired seeding or impaired outgrowth, we next aimed to determine which step in metastasis is dependent on E-selectin.

To delineate the role of E-selectin in metastatic seeding, we evaluated the livers of wild-type or E-selectin KO mice one day following splenic injection of CA19-9^pos^ KPCY PDAC cells (Figure 3I). We found significantly reduced numbers of YFP^+^ cells present in the livers of E-selectin KO mice (Figure 3J). Similarly, neutralization of CA19-9 or E-selectin using antibody pre-treatment significantly suppressed metastatic seeding (Figure 3K-M). These data suggest that both CA19-9 and E-selectin promote seeding of liver metastasis.

Together, these findings demonstrate that CA19-9 enhances the initial seeding of liver metastases *in vivo* through interaction with E-selectin on the vasculature, establishing CA19-9 as a functional mediator of metastasis.

### CA19-9-mediated liver metastasis is not associated with major changes in immune cell infiltration

Aberrant glycosylation has been reported to promote tumor progression and metastasis through immune evasion (9, 37–40). To determine whether immune-mediated mechanisms contribute to the enhanced metastatic burden observed in CA19-9^pos^ tumors, we assessed liver metastasis in immunocompromised NSG mice. Despite the absence of adaptive immunity, CA19-9^pos^ cells still produced markedly larger liver metastases than CA19-9^neg^ cells (Supplemental Figure 7A-E). Additionally, we examined whether CA19-9 expression altered immune cell infiltration within liver metastases in immunocompetent WT mice. IHC quantification of CD8⁺ T cells, Foxp3⁺ regulatory T cells, and F4/80⁺ macrophages revealed no significant differences between CA19-9^neg^ and CA19-9^pos^ liver metastasis (Supplemental Figure 7F-H), indicating that CA19-9 does not substantially impact the abundance of these major immune cell subsets in metastatic lesions. These findings suggest that the effect of CA19-9 on liver metastasis is unlikely to be driven by changes in immune cell infiltration.

### CA19-9 promotes outgrowth of liver metastasis

To evaluate whether therapeutic targeting of CA19-9 can impact liver metastasis outgrowth independent of seeding, we treated mice bearing CA19-9^pos^ PDAC tumors with the anti-CA19-9 antibody (5B1) or isotype control (ISO) three days after splenic injection (Figure 4A). CA19-9 blockade resulted in a striking reduction in liver weight (Figure 4B-D) as well as CK19 positive tumor area compared with isotype-treated controls (Figure 4E, F). The number and diameter of metastatic foci were also significantly decreased (Figure 4G, H), indicating that CA19-9 blockade suppresses the outgrowth of liver metastasis. While proliferation was not significantly different at endpoint (Figure 4I), the number of apoptotic cells was significantly increased in CA19-9 antibody-treated tumors (Figure 4J).

**Figure 4.**
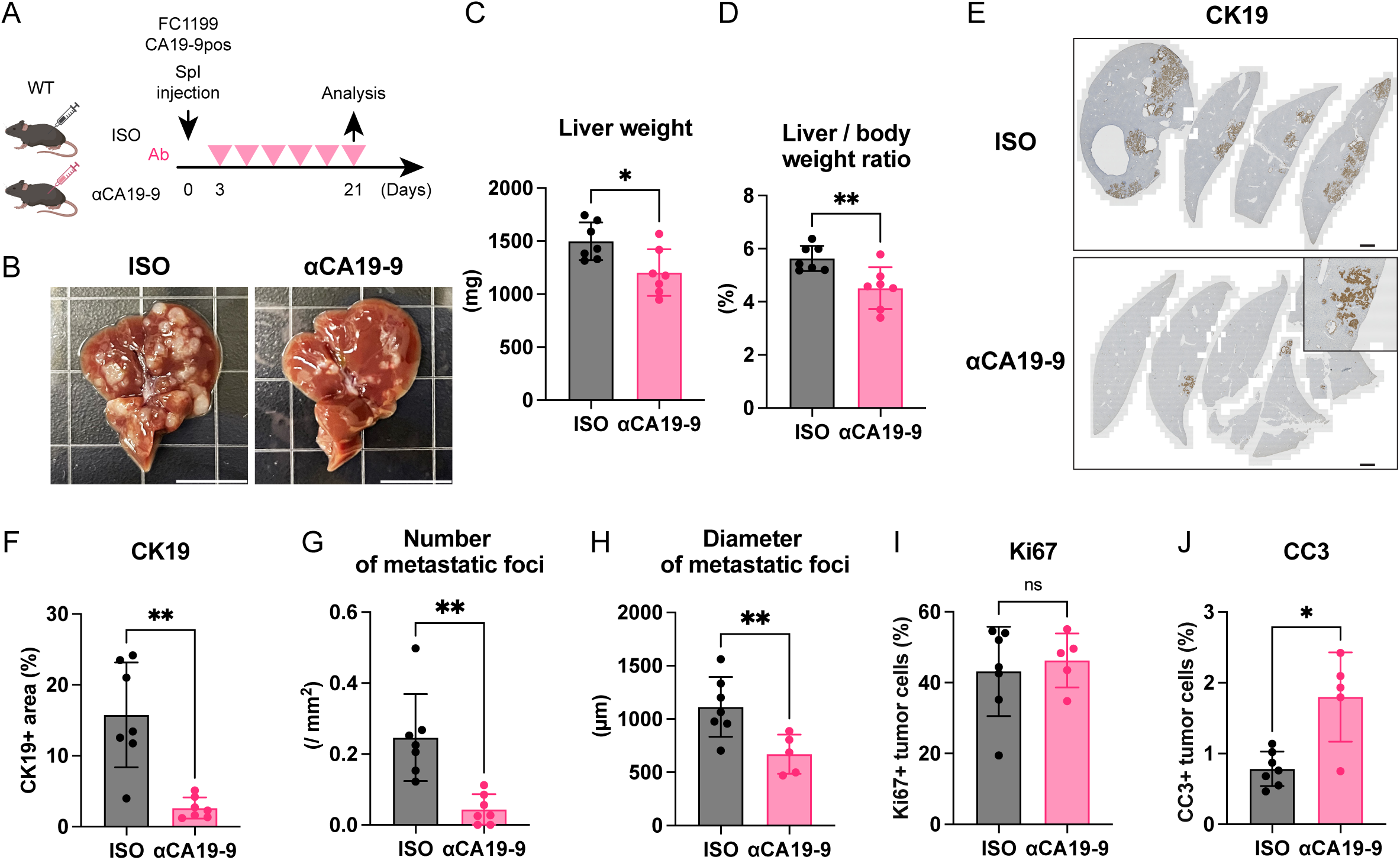
Anti-CA19-9 antibody suppresses the liver metastasis *in vivo*. (A) Schema of study design in (B-J) (B) Representative macroscopic images of livers isolated from mice injected with CA19-9^pos^ cells and treated with either isotype control (ISO) (left) or anti-CA19-9 antibody (5B1; αCA19-9) (right). Scale bar = 1 cm. (C, D) Quantification of liver weight (C) and of liver weight normalized by body weight (D). (E) Representative CK19 IHC of whole liver sections in splenic injection models treated either isotype control (ISO) or anti-CA19-9 antibody (5B1; αCA19-9). Scale bar = 1 mm. (F) Quantification of CK19 positive area (%) across the whole liver sections. (G) Number of metastatic foci across the whole liver sections normalized by tissue area (mm²). (H) Mean diameter of metastatic foci measured across the whole liver sections. (I-J) Quantification of Ki67 positive (I) and cleaved caspase-3 (CC3) positive (J) tumor cells within liver metastasis area. *Data are presented as mean ± SD. Each dot represents an individual mouse. Mice were injected with 1 × 10^5^ tumor cells per mouse. Sample sizes may vary between analyses because some liver sections lacked detectable metastatic lesions. Statistical significance was calculated using unpaired two-tailed t-test with Welch’s correction. *P < 0.05; **P < 0.01; ns, not significant.

To investigate the extent to which E-selectin plays a role in the metastatic cascade post-extravasation and seeding, we implemented a post-operative E-selectin treatment regimen (Supplemental Figure 8A). By initiating anti-E-selectin antibody administration after splenic injection, we were able to assess whether E-selectin influences the outgrowth of established metastatic foci rather than the initial seeding process. Treatment with the anti-E-selectin antibody three days after surgery did not suppress the metastatic burden compared with isotype control (Supplemental Figure 8B-D). These data indicate that E-selectin primarily mediates cancer cell extravasation and seeding rather than outgrowth.

Together, these results indicate that while E-selectin is primarily involved in metastatic seeding, CA19-9 plays a role in both seeding and outgrowth of PDAC metastases. These data suggest that there is a large window for anti-CA19-9 targeted therapy for the treatment of metastatic PDA. We next sought to delineate the mechanisms by which CA19-9 promotes PDAC metastatic outgrowth.

### CA19-9 expression activates the AKT signaling pathway in pancreatic cancer cells and liver metastases

To investigate whether CA19-9 modulates intracellular signaling pathways, we performed RNA-seq analysis on CA19-9^pos^ and CA19-9^neg^ FC1199 and FC1245 cells. Gene set enrichment analysis (GSEA) demonstrated significant enrichment of the PI3K-AKT-mTOR and mTORC1 hallmark signatures in CA19-9^pos^ cells (Figure 5A). We validated these findings at the protein level. CA19-9^pos^ FC1199 cells exhibited increased phosphorylation of AKT and S6 compared with CA19-9^neg^ cells (Figure 5B, C). In contrast, phosphorylation of ERK1/2 was not upregulated, suggesting that CA19-9 enhances AKT but not MAPK signaling.

**Figure 5.**
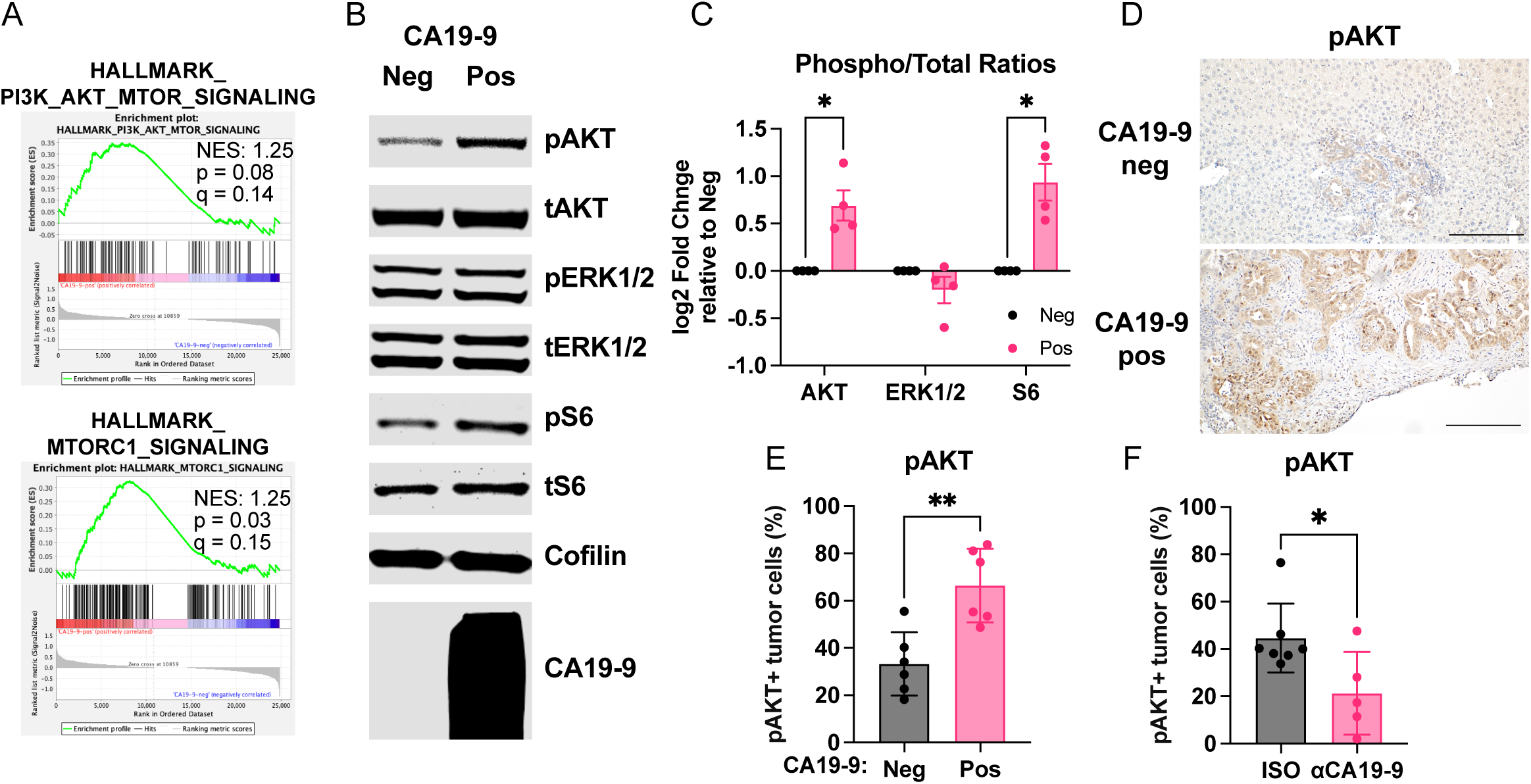
CA19-9 activates pAKT signaling pathway in pancreatic cancer cells and liver metastasis. (A) Gene set enrichment analysis (GSEA) of RNA-seq data comparing CA19-9^neg^ and CA19-9^pos^ FC1199 cells. Hallmark PI3K-AKT-mTOR signaling (top) and mTORC1 signaling (bottom) pathways were enriched in CA19-9^pos^ cells. (B) Representative immunoblot images of CA19-9^neg^ and CA19-9^pos^ FC1199 cells cultured in low-serum conditions for 24 hrs. (C) Quantification of phospho/total protein ratios for AKT, ERK, and S6 shown in (B). (D) Representative pAKT IHC in liver metastases derived from CA19-9^neg^ or CA19-9^pos^ FC1199 cells. Scale bar = 200 μm. (E) Quantification of pAKT positive tumor cells in liver metastasis derived from CA19-9^neg^ or CA19-9^pos^ FC1199 cells. (F) Quantification of pAKT positive tumor cells in liver metastasis of CA19-9^pos^ FC1199 cells in mice treated with either isotype control (ISO) or anti-CA19-9 antibody (5B1; αCA19-9). *Data are presented as mean ± SD. (C) Each dot represents one independent experiment. (E, F) Each dot represents an individual mouse. Statistical significance was calculated using unpaired two-tailed t-test with Welch’s correction. *P < 0.05; **P < 0.01.

We next examined whether AKT phosphorylation is increased in CA19-9^pos^ liver metastasis *in vivo*. pAKT IHC showed significantly higher expression in CA19-9^pos^ than in CA19-9^neg^ liver metastasis, while pERK1/2 remained unchanged (Figure 5D, E, and Supplemental Figure 9A-B). In addition, pAKT levels were significantly reduced following CA19-9 blockade (Figure 5F). These results indicate that CA19-9 promotes activation of AKT-mTOR signaling both *in vitro* and *in vivo*, suggesting that the increased outgrowth of liver metastasis observed in CA19-9^pos^ tumors may be AKT dependent.

To place these findings in a clinical context, we performed immunofluorescence staining for pAKT and CA19-9 using a tissue microarray (TMA) of primary PDAC tissues to examine the association between CA19-9 expression and AKT phosphorylation in human PDAC. Quantitative image analysis revealed a positive correlation between the percentage of CA19-9-positive tumor cells and pAKT-positive tumor cells (Figure 6A, B). Moreover, CA19-9-high tumors displayed significantly increased levels of pAKT-positive tumor cells compared with CA19-9-low tumors (Figure 6C). To investigate the effect of CA19-9 on the prognosis of PDAC patients while minimizing potential confounding by tumor burden, we re-analyzed a published PDAC cohort and focused our analysis on patients with stage IA disease (T1N0M0; tumor size ≤ 2 cm, R0) (41). Notably, even within this early-stage cohort, elevated preoperative CA19-9 levels were associated with significantly worse disease-free survival (DFS) and overall survival (OS) (Figure 6D, E), suggesting that CA19-9 may reflect biological properties not fully captured by tumor size alone.

**Figure 6.**
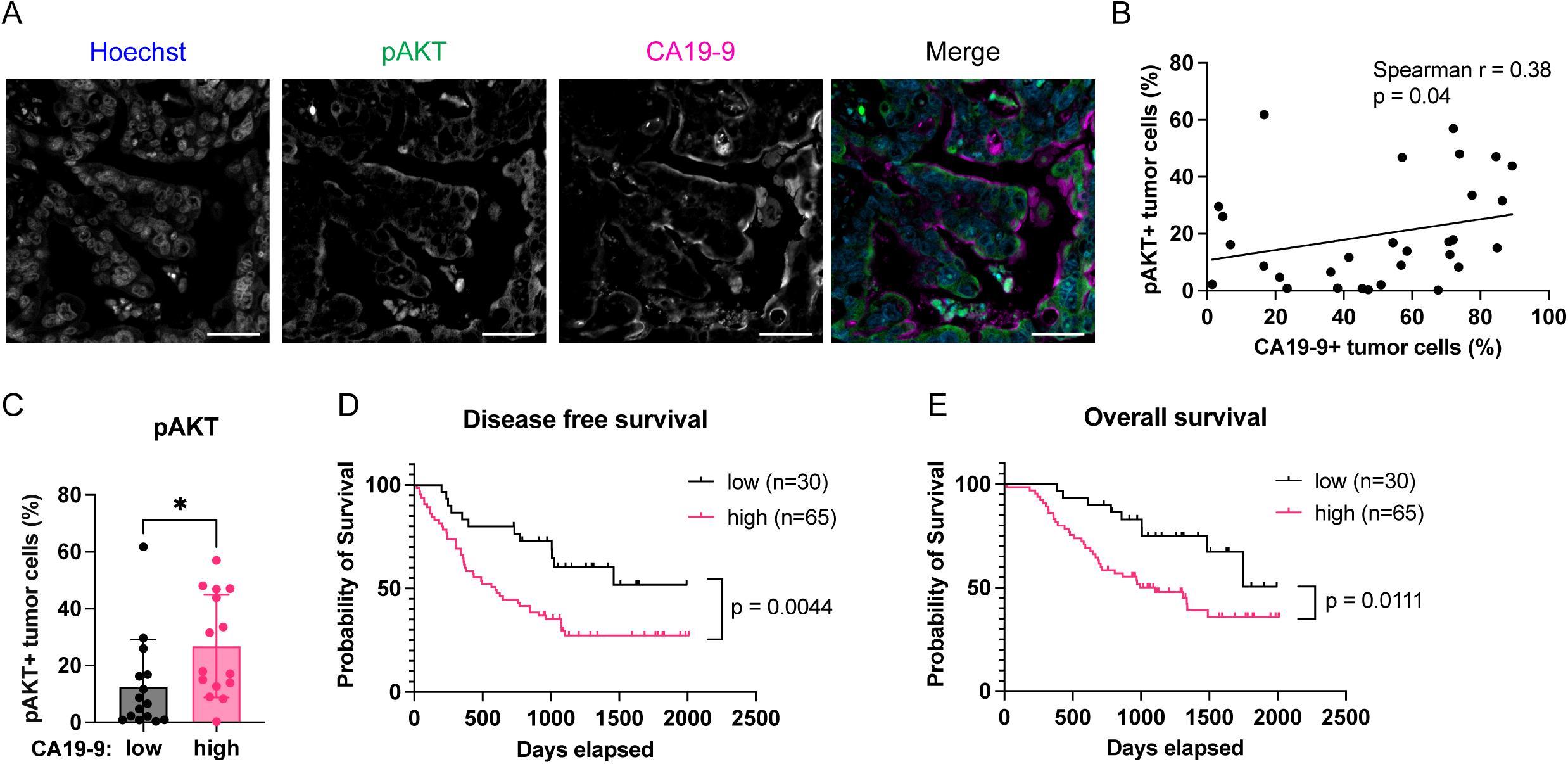
CA19-9 expression is associated with AKT activation and poor clinical outcomes in patients with PDAC. (A) Representative immunofluorescence images of TMA stained for Hoechst (blue), phosphorylated AKT (pAKT; green), and CA19-9 (magenta). The merged image is shown on the right. Scale bar = 50 μm. (B) Correlation between the percentage of CA19-9-positive tumor cells and pAKT-positive tumor cells. Spearman correlation analysis was performed to evaluate the association. The solid line indicates the linear regression fit. (C) Comparison of pAKT-positive tumor cells between CA19-9-low and CA19-9-high tumors. Tumors were classified based on the median percentage of CA19-9-positive tumor cells. Statistical significance was calculated using the Mann-Whitney *U* test. (D, E) Kaplan-Meier analyses of disease-free survival (DFS) (D) and overall survival (OS) (E) in patients with stage IA PDAC, stratified by preoperative serum CA19-9 levels (CA19-9-low, ≤37 U/mL; CA19-9-high, >37 U/mL). For DFS, the hazard ratio (HR) for CA19-9-low versus high tumors was 0.46 (95% CI, 0.27-0.79). For OS, the HR was 0.46 (95% CI, 0.25-0.84). Statistical significance was calculated using the log-rank (Mantel-Cox) test. HRs and 95% confidence intervals (CIs) were derived using the Mantel-Haenszel method. *Data are presented as mean ± SD. Each dot represents an individual patient. *P < 0.05.

## Discussion

CA19-9 is widely used in clinical practice as a biomarker for disease monitoring in patients with PDAC (13). Despite its strong association with poor prognosis and early postoperative recurrence in multiple clinical studies (15–20), CA19-9 has largely been regarded as a passive marker, and its functional contribution to metastasis has remained unclear. Although interactions between CA19-9 and E-selectin have been well documented *in vitro*, the *in vivo* relevance of this axis has been difficult to assess due to the lack of endogenous CA19-9 expression in mice. In this study, we overcame this limitation by establishing syngeneic CA19-9-positive murine PDAC models and demonstrated that CA19-9 actively promotes liver metastasis through multiple steps of the metastatic cascade. Our data support a functional role of CA19-9 in liver metastasis: facilitating initial seeding via E-selectin-mediated adhesion and enhancing metastatic outgrowth through activation of AKT signaling. These findings provide *in vivo* evidence that CA19-9 is not merely a biomarker, but a functional mediator of metastatic progression in PDAC.

Although glycosylated proteins can contribute to immune evasion (9, 37–40), our analyses indicated minimal differences in immune cell infiltration between CA19-9 negative and positive metastases. Instead, CA19-9-positive tumor cells exhibited elevated AKT signaling. AKT activation has been shown to enhance tumor cell survival and colonization and to promote metastatic outgrowth in the liver (42). This suggests that CA19-9 may enhance the survival of metastatic cells rather than directly affecting tumor cell proliferation. Additionally, recent spatial transcriptomic analyses have shown upregulation of AKT signaling in liver metastases compared with primary pancreatic tumors (43), implying that AKT activation confers a selective advantage within the hepatic niche. Moreover, CA19-9 blockade reduced pAKT levels and suppressed CA19-9-positive metastatic outgrowth in the liver, suggesting CA19-9 can be a therapeutic target at multiple stages of tumor progression.

Prior work indicates that E-selectin blockade can disrupt tumor-endothelial adhesion in acute myeloid leukemia and lung cancer (44, 45), consistent with our findings that CA19-9 promotes hepatic seeding of PDAC cells through E-selectin-mediated interactions. Strategies targeting the CA19-9-E-selectin axis, using the humanized monoclonal anti-CA19-9 antibody (clone 5B1, also known as BNT321) or pharmacologic inhibition of E-selectin have been reported to be feasible in clinical trials (27, 28). Our data indicate that E-selectin-directed approaches may preferentially modulate early adhesive interactions during metastatic seeding, whereas direct targeting of CA19-9 has the potential to interfere with multiple stages of metastatic progression.

Regarding the clinical translation of the anti-CA19-9 antibody, it has been reported that antibody preloading improves antibody accumulation within tissues to mitigate binding to serum CA19-9 (46), indicating that further optimization of dosing strategies may enhance clinical efficacy. This implies that anti-CA19-9 antibody may be most effective in the postoperative adjuvant setting, where the tumor burden is minimal and circulating antigen levels are lower. The clinical safety of anti-CA19-9 antibody in a phase 1 study has already been reported (27). Although mFOLFIRINOX+BNT321 Phase 1/1b trials as adjuvant therapy following curative resection (NCT06069778) or in metastatic, advanced line disease (NCT02672917) were terminated by sponsor decision, initial findings in the metastatic setting reported stable disease in 20% of monotherapy treated patients (>90 days) and partial response was observed in 27% of patients who received the antibody with chemotherapy (>10 months on study) (27). Together, these observations suggest that CA19-9 targeted therapies may be effective when deployed in multiple settings.

Our analyses were primarily focused on liver metastasis, which represents the most common metastatic site. Given its role in promoting tumor cell seeding in the liver, it is plausible that CA19-9 may also facilitate metastatic dissemination to other vascular beds. However, whether CA19-9 similarly contributes to metastatic progression in other organs remains to be determined. While E-selectin emerged as a key mediator of CA19-9 dependent hepatic seeding, additional CA19-9 interacting lectins or adhesion molecules may also contribute to metastatic progression. Future studies dissecting the contribution of specific CA19-9 modified glycoproteins that facilitate liver metastasis through cell intrinsic mechanisms or through interactions within the hepatic microenvironment are needed to reveal additional therapeutic interception points.

Overall, our findings reveal that CA19-9 plays an important role in promoting the seeding and outgrowth of liver metastasis, providing a mechanistic rationale for its strong clinical association with postoperative recurrence and highlighting a promising therapeutic target in pancreatic cancer.

## Methods

### Sex as a biological variable

This study included experiments with human samples and animal models. The human samples included both males and females. Male and female mice were used in this study. To match the biological sex of the tumor cell lines, male-derived cell lines were implanted into male mice and female-derived cell lines into female mice. Similar findings are reported for both sexes.

### Animal models

C57BL/6J (Jackson Laboratory Strain #: 000664) and E-selectin KO (Jackson Laboratory Strain #: 008236) mice were maintained in a pathogen-free animal facility at the Salk Institute for Biological Studies. Animals were housed with a 12-h light-dark cycle, at an ambient temperature (22O±O1O°C), and 30-70% humidity. Water and food were provided ad libitum.

Both male and female mice between 8 and 14Oweeks of age were used. In all experiments, age-matched and sex-matched mice were randomly assigned to experimental groups. Therapeutic antibody intervention studies were performed using intraperitoneal injection of 200 μg of antibody twice a week. For CA19-9 targeting, mice received either anti-CA19-9 antibody (5B1; MabVax (acquired by BioNTech)) or a human IgG isotype control (BioXcell, BE0297). For E-selectin targeting, mice received either anti-E-selectin antibody (9A9; BioXcell, BE0294) or a rat IgG isotype control (Leinco Technologies, Inc., I-1034).

### Cell lines

The mouse PDAC cell lines (FC1199, male and FC1245, female) were derived from Pdx1-Cre; Kras^LSL-G12D/+^; Trp53^LSL-R172H/+^ (KPC) mice (C57BL/6J) and provided by Dr. David Tuveson (Cold Spring Harbor Laboratory). The KPCY mouse pancreatic cancer cell line 6419c5 (female) was obtained from Kerafast. CA19-9 expressing cells were generated as previously described (26). SW1990 (ATCC CRL-2172) and Capan-2 (ATCC HTB-80) were originally obtained from the American Type Culture Collection (ATCC). All cells were maintained in DMEM (Dulbecco’s Modified Eagle Medium) supplemented with 10% fetal bovine serum (FBS) and 1% penicillin/streptomycin (P/S) at 37°C with 5% CO2. The hM19a 2D cell line was generated from hM19a human pancreatic cancer organoid established at Cold Spring Harbor Laboratory (29). The organoids were directly seeded onto standard six-well tissue culture plates without extracellular matrix support and maintained in DMEM supplemented with 20% FBS and 1% P/S. Human Umbilical Vein Endothelial Cells (HUVECs) (ScienceCell Catalog No.#8000) were maintained in Endothelial Cell Medium (ScienceCell Catalog No.#1001) supplemented with ECGS, FBS and P/S, following the manufacturer’s instructions. All cell lines underwent routine mycoplasma testing.

### Splenic injection model (liver metastasis model)

For the experimental splenic injection model, mice were anesthetized under continuous isoflurane and administered sustained-release buprenorphine. A small left flank incision was made to expose the spleen. After ligation of the center of the spleen in two places between splenic blood vessels at both splenic poles, the spleen was divided between the ligation clips and the upper pole of the spleen was placed into the peritoneum. Murine PDAC cells (5 × 10^4^, 1 × 10^5^ or 3 × 10^5^ per mouse), suspended in 50 μL of medium, were injected into the lower pole of the spleen using a 30G insulin syringe. The number of injected cells was adjusted depending on the experimental design, as indicated in the corresponding figure legends. The injected pole of the spleen was removed by ligating splenic vessels at the distal end of the pancreas after 1-2 minutes to allow cell dissemination. The incision was closed with absorbable sutures and wound clips. Mice were euthanized at timed endpoints for analysis of liver metastases. Liver tissues were collected, fixed in 10% neutral-buffered formalin, and processed for histopathology.

### Cell viability assay (CellTiter-Glo)

Cell growth was measured using the CellTiter-Glo Luminescent Cell Viability Assay (Promega) following the manufacturer’s instructions. Briefly, cells were seeded into white-walled 96-well plates at the density of 3,000 cells per well in 100 μL of medium. An equal volume of CellTiter-Glo reagent was added to each well at Day 0, 1, 3 and 5. Luminescence was measured using a microplate reader (Tecan Spark Cyto) to quantify ATP levels as an indicator of metabolically active cells. All measurements were performed in triplicate.

### Scratch (Wound healing) assay

Cell migration was evaluated using a scratch assay monitored in real time using IncuCyte live-cell imaging system (Sartorius). Cells were seeded into 96-well ImageLock plates (Essen BioScience) in three or more replicates and grown to full confluence. A uniform scratch was created using the IncuCyte WoundMaker tool, followed by washing with PBS to remove detached cells. Serum-free medium was added, and images were captured every 4 hours for up to 72 hours using the IncuCyte system. Wound closure was quantified using IncuCyte software.

### Adhesion assay

Near-confluent HUVECs cultured in 96-well plates were stimulated with TNF-α (10 ng/mL) for 6-8 hours to induce E-selectin expression as previously described (21–23). Tumor cells were labeled with Calcein-AM (Cayman Chemical 400146-1) for 30 minutes at 37°C. The labeled tumor cells (5 × 10^4^ cells per well) were added to stimulated HUVECs in three replicates and incubated for 30 minutes at room temperature on a tilting shaker to minimize nonspecific binding. Nonadherent cells were removed by gently washing with PBS three times. Adherent cells were imaged using a TECAN Spark system and quantified by counting Calcein-positive cells per field in QuPath version 0.5.1. For antibody blocking experiments, tumor cells were incubated with human IgG (BioXcell, BE0297) or anti-CA19-9 antibody (5B1; MabVax (acquired by BioNTech)) at 50 µg/mL for 30 minutes at 37°C with gentle rotation. Alternatively, HUVECs were incubated with mouse IgG (Leinco Technologies, Inc., I-536) or anti-human E-selectin Ab (R&D Systems, BBA16) at 25 µg/mL for 30 minutes at 37°C.

### Generation of FUT3 knockout cells using CRISPR/Cas9 system

FUT3-knockout Capan-2 cells were generated using a lentiviral CRISPR/Cas9 system. Capan-2 cells were first transduced with a lentiviral vector encoding Cas9 and selected with blasticidin to establish stable Cas9-expressing cells. Single-guide RNAs (sgRNAs) targeting FUT3 did not achieve sufficient knockout efficiency, so three independent sgRNAs targeting FUT3 were co-introduced. The sgRNA sequences were as follows: sgFUT3-1, TCCCAGTGGGTCCTCCCGAC; sgFUT3-2, CGATGCCACTGGATCCCCTA; and sgFUT3-3, CAGCGGCGCCATGGCCATTG. As a negative control, we used a control sgRNA targeting a noncoding genomic locus. Transduced cells were selected with puromycin. After antibiotic selection, cells were stained with NS19-9 antibody followed by anti-mouse Alexa Fluor 488 secondary antibody, and the CA19-9-negative population was sorted by flow cytometry. Loss of CA19-9 was confirmed by immunoblot analysis.

### Immunohistochemistry and immunofluorescence

Tissues were fixed in 10% neutral-buffered formalin, embedded in paraffin, and sectioned at 5 μm thickness. For immunohistochemistry (IHC), heat-induced antigen retrieval was performed in citric acid buffer (pH 6.0), EDTA buffer (pH 8.0), or EDTA/Tris buffer (pH 9.0) in a pressure cooker for 20 minutes after dewaxing. Endogenous peroxidases were quenched with 3% H2O2 in Tris-buffered saline with 0.1% Tween20 (TBST) for 15 minutes at room temperature. Tissue sections were blocked with 10% goat serum and 2.5% horse serum. For experiments involving goat-derived primary antibodies, blocking was performed using 2.5% horse serum. Then, sections were incubated with primary antibody overnight at 4°C. Antibodies are described in Supplemental Table S1. The sections were incubated with secondary antibody (Vector Laboratories, ImmPRESS HRP IgG Polymer Detection Kit, MP-7451, MP-7404, MP-7405, PI-3000) for 30 minutes at room temperature after washing with TBST. The signals were developed using ImmPACT DAB HRP Substrate kit (Vector Laboratories, SK-4105) after washing with TBST. Slides were counterstained with hematoxylin, dehydrated, and coverslipped. Slides were imaged using an Olympus BX43 Microscope or a Zeiss Axioscan 7. CellSens or Zen lite was used to visualize and annotate images.

For immunofluorescence (IF), after antigen retrieval, tissue sections were permeabilized with 0.5% Triton X-100 in PBS for 10 minutes and blocked with 2.5% normal horse serum for 1 hour at room temperature. Sections were then incubated with primary antibodies overnight at 4°C. Antibodies are described in Supplemental Table S1. After washing with TBST, sections were incubated with Alexa Fluor-conjugated secondary antibodies (Life Technology A-31572, Jackson ImmunoResearch #715-605-151) for 1 hour at room temperature in the dark, followed by counterstaining with Hoechst (Thermo Fisher Scientific). Slides were mounted with ProLong Glass Antifade Mountant (Thermo Fisher Scientific P36980) and imaged using a Zeiss Axioscan 7. Zen lite was used to visualize and annotate images.

### Quantification of IHC staining

IHC slides were imaged and analyzed in QuPath version 0.5.1. DAB-positive cells or areas were quantified using brightfield (H-DAB) settings after stain vector estimation. For CK19 staining, whole-slide images were scanned using a Zeiss Axioscan 7 and analyzed to assess total tumor burden. The CK19-positive area was normalized to the total hematoxylin-positive tissue area to calculate the CK19-positive area fraction per section. Liver tissues were embedded to include multiple lobes and diverse anatomical regions within a single paraffin block, thereby minimizing sampling bias across metastatic sites. One representative section per mouse was analyzed, and each data point represents one mouse. For other markers, 3-5 metastatic regions per section were randomly selected when available and imaged using an Olympus BX43 microscope.

After annotation of tumor areas, tumor cells were identified and classified using QuPath’s cell detection and object classification tools based on nuclear detection and morphological features. The percentage of marker-positive tumor cells was calculated using a predefined DAB intensity threshold that was uniformly applied. Positive tumor cells were normalized to the total number of classified tumor cells within the annotated regions. The mean value per mouse was used for statistical analysis. Each data point represents one mouse.

### Quantification of tissue microarray

A commercial tissue microarray (TMA) containing duplicate cores from 40 human PDAC cases was purchased from US Biomax (PA1002b). Sections were stained for CA19-9 and pAKT by immunofluorescence and imaged using a Zeiss Axioscan 7. Images were analyzed using QuPath (version 0.5.1). After TMA dearraying, cells were detected based on nuclear staining with Hoechst. Tumor cells were identified using QuPath’s cell detection and object classification tools based on nuclear detection and morphological features. Cores containing staining artifacts or few tumor cells were excluded from analysis. The percentages of CA19-9-positive tumor cells and pAKT-positive tumor cells among total tumor cells were calculated for each core. The two cores corresponding to each case were analyzed independently, and the mean value was used for correlation analysis. Spearman correlation analysis was performed to evaluate the association between CA19-9 positivity and pAKT positivity across cases.

### Immunoblotting

Cells were lysed in TNET (1% (v/v) Triton X-100, 150 mM NaCl, 5 mM EDTA, 50 mM Tris-HCl pH 7.5) or RIPA buffer supplemented with 1x protease and phosphatase inhibitors (Life Technologies #A32961). Lysates were incubated on ice for 15 minutes and clarified by centrifugation at 10,000 g for 10 minutes at 4°C. Protein concentration was measured using a BCA assay (Thermo Fisher Scientific #23227). For immunoblotting, lysates were mixed with NuPAGE™ LDS Sample Buffer (4X) (Thermo Fisher Scientific # NP0007) and 10X Bolt Sample Reducing Agent (Thermo Fisher Scientific #B0009), heated to 95°C for 10 minutes, separated on 4-12% Bis-Tris NuPAGE gels (Thermo Fisher Scientific NP0321BOX), and then transferred to PVDF membranes (Thermo Fisher Scientific # 88518 for ECL, MilliporeSigma IPFL00010 for fluorescence). After blocking in 5% BSA for chemiluminescent detection or 5% milk for fluorescence-based detection for one hour, membranes were incubated with primary antibodies (Supplemental Table S1) overnight at 4°C with gentle rocking. After incubation with HRP-conjugated secondary antibodies (Jackson ImmunoResearch #115-035-003, #711-035-152, #109-035-003) for 2 hours, signals were detected using enhanced chemiluminescence (Li-COR 926-95000) and imaged using the Odyssey M in chemiluminescence mode or autoradiography. Alternatively, membranes were incubated with fluorophore-conjugated secondary antibodies (Li-COR #926-32211, #926-68070) for 2 hours. The signals were visualized using the Odyssey M (Li-COR). Quantification was performed using Empiria Studio (Li-COR).

### E-selectin pull-down assay

FC1199 CA19-9 negative and positive murine PDAC cells were cultured in serum-free DMEM for 72 hours to generate conditioned media. Supernatants were collected and concentrated by centrifugation at 4,000 g for 2 hours. Pull-down assays were performed using Dynabeads Protein G for Immunoprecipitation (Invitrogen, 10004D) in combination with recombinant mouse E-selectin/CD62E Fc Chimera Protein (Fisher scientific #724ES100) or recombinant human E-selectin/CD62E Fc Chimera Protein (Fisher Scientific #575ES100). Procedures were conducted according to the manufacturer’s protocol with the following modifications: wash and binding steps were performed using 20 mM HEPES, 150 mM NaCl (pH 7.5), and antigen incubation was carried out overnight at 4°C with rotation. Following immunoprecipitation, bound proteins were eluted in NuPAGE™ LDS Sample Buffer (4X) and 10X Bolt Sample Reducing Agent, boiled at 95°C for 15 minutes, and subjected to SDS-PAGE. Immunoblotting was performed as described above. Membranes were probed with NS19-9 (1 µg/mL) and signals were detected by enhanced chemiluminescence and visualized using autoradiography.

### RNA isolation and RNA-seq

Cells were lysed directly with 1 mL of TRIzol reagent (Thermo Fisher Scientific #15596018), and total RNA was extracted using RNeasy Mini Kit (Qiagen #74104) according to the manufacturer’s instructions. RNA concentration was measured using a NanoDrop One (Thermo Fisher Scientific). RNA quality and integrity were assessed using a Qubit Fluorometer (Invitrogen) and a TapeStation (Agilent) before RNA-seq analyses. RNA-seq libraries were prepared from total RNA using an rRNA depletion protocol and sequenced at the Integrative Genomics Core of the Salk Institute on a NovaSeq 6000 platform (Illumina) with 50-bp paired-end reads. For FC1199 cells, three biological replicates were analyzed for each group (CA19-9^neg^ and CA19-9^pos^). For FC1245 cells, three CA19-9^neg^ and two CA19-9^pos^ samples passed quality control and were included in the final analysis.

### Bulk RNA-seq analysis

Raw reads were quality-checked with FastQC v0.11.8 (30). Experiments containing adaptor sequences were trimmed with Trim Galore v0.4.4_dev (31). Reads were aligned to the mm10 reference genome with STAR aligner v2.5.3a (32), and converted to gene counts with HOMER’s analyzeRepeats.pl script (33). Gene counts were normalized and queried for differential expression using DESeq2 v1.30.0 (34). Gene set enrichment analysis (GSEA) was performed using the GSEA software (Broad Institute). Enrichment analysis was conducted using the Hallmark gene sets from the Molecular Signatures Database (MSigDB). Normalized enrichment scores (NES) and FDR q values were used to assess statistical significance.

### Statistics

Data are presented as mean ± SD unless otherwise indicated. Statistical analyses were performed using GraphPad Prism version 11 (GraphPad Software). Normality was assessed using the Shapiro-Wilk test. For 2-group comparisons, either the 2-tailed unpaired t-test with Welch’s correction or the Mann-Whitney *U* test was used as appropriate. Correlations were assessed using Spearman’s rank correlation coefficient. Survival estimates were generated using the Kaplan-Meier method and compared using the log-rank test. HRs and 95% confidence intervals (CIs) were derived using the Mantel-Haenszel method. The specific statistical tests used for each analysis are indicated in the figure legends. A P value less than 0.05 was considered statistically significant.

### Study approval

All animal experiments were performed in accordance with protocols approved by the Institutional Animal Care and Use Committee at the Salk Institute for Biological Studies (Protocol #18-00048).

### Data availability

All data associated with this study are available in the article or the Supporting Data Values file. RNA-seq data generated in this study from murine PDAC cells will be deposited in the Gene Expression Omnibus (GEO) upon publication.

## Author contributions

SO and DDE designed research studies. SO, HS, JH, VP, VOV, CSK, JRT, CRB, KLP, YW, KC, SK and RT conducted experiments and acquired data. SO, KL, JZhu and JZou analyzed data. MD, RME, AML and HT provided reagents and materials. SO and DDE wrote the manuscript with input from all authors.

## Acknowledgements

KPCY 6419c5 cells were kindly provided by Dr. Susan Kaech in the Salk Institute. SW1990 cells were kindly provided by Dr. Yuan Sui and Dr. Tony Hunter in the Salk Institute. HUVECs were kindly provided by Christina K. Lim and Dr. Fred H Gage in the Salk institute. The authors would like to thank all the members and alumni of the Engle laboratory, the Animal Resources Department (ARD) of the Salk Institute, the UCSD Biorepository and Tissue Technology Shared Resource (BTTSR), and Daniela Boassa, Sammy Weiser Novak and Elsie Quansah at the Biophotonics Core Facility of the Salk Institute.

