## Supplemental Figures and Tables for "CA19-9 promotes liver metastasis of pancreatic cancer through E-selectin mediated extravasation"

Supplemental Figure 1

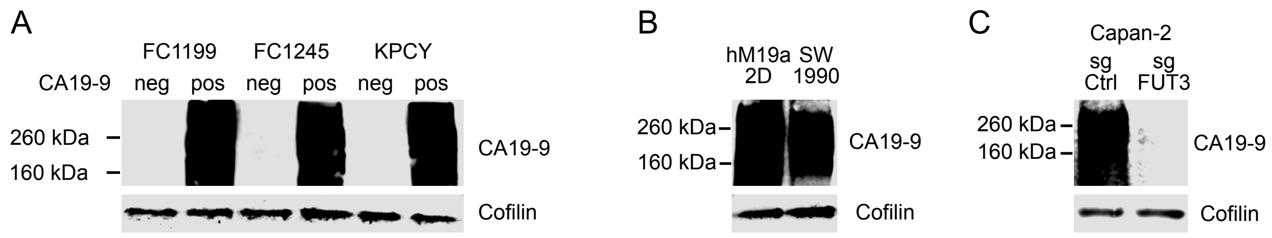


**Supplemental Figure 1. CA19-9 expression in mouse and human PDAC cell lines.**
(A) Immunoblot analysis of CA19-9 expression in CA19-9neg and CA19-9pos murine PDAC cells (FC1199, FC1245, and KPCY).
(B) Immunoblot analysis of CA19-9 expression in human CA19-9pos cell lines (hM19a 2D and SW1990). CA19-9 was detected using the anti-CA19-9 antibody 5B1.
(C) Immunoblot analysis of CA19-9 expression in Capan-2 cells with control sgRNA (sgCtrl) or FUT3-targeting sgRNA (sgFUT3). CA19-9 was detected using the anti-CA19-9 antibody 5B1.

Supplemental Figure 2

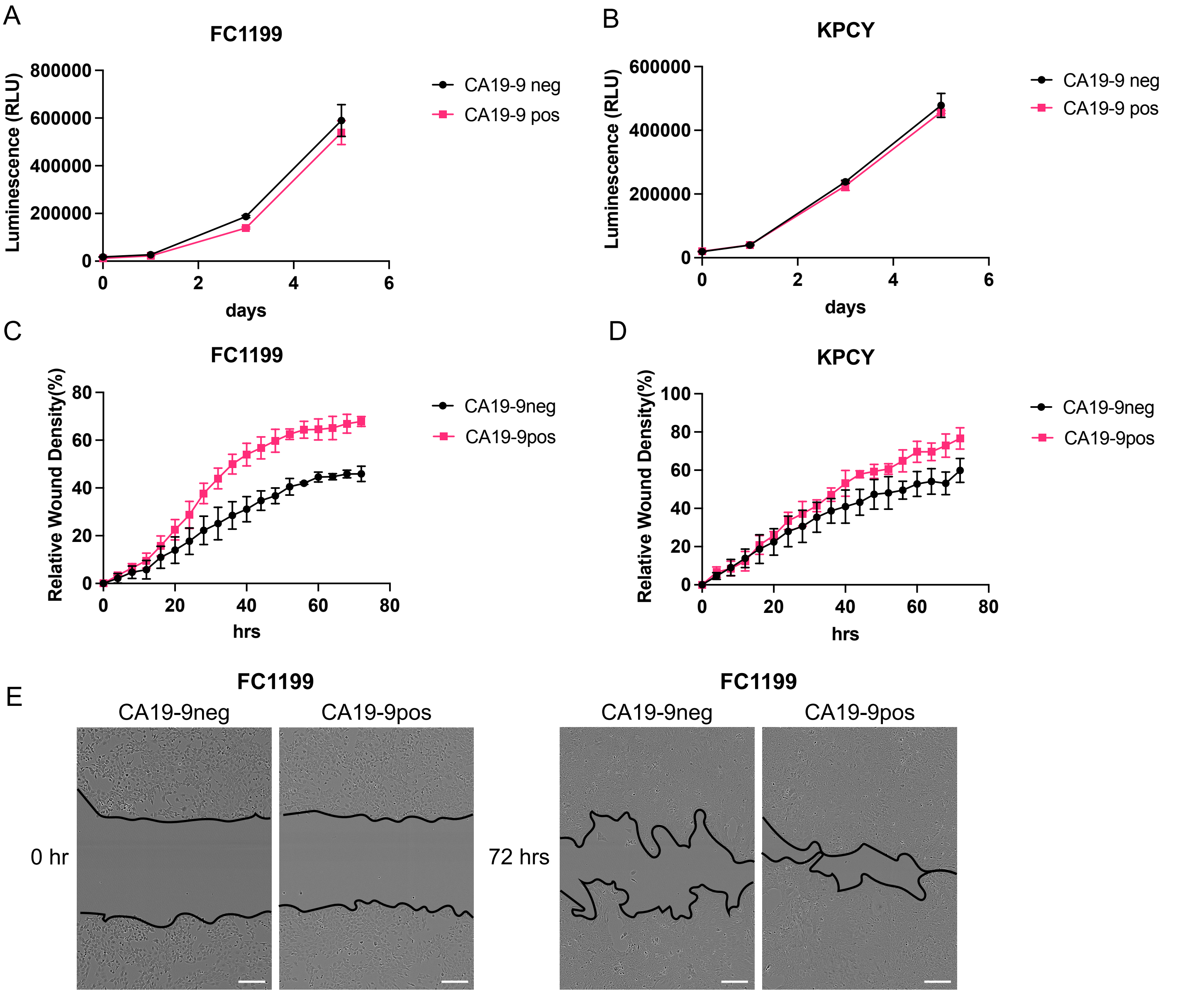


**Supplemental Figure 2. CA19-9 expression does not alter proliferation but enhances migratory capacity in vitro.**

(A, B) Cell viability assay of CA19-9neg and CA19-9pos FC1199 (A) and KPCY (B) cells.

(C, D) Scratch wound healing assay of CA19-9neg and CA19-9pos FC1199 (C) and KPCY (D) cells. Wound closure was normalized to the wound area at time 0 and relative wound density was quantified.

(E) Representative images of Scratch wound healing assay using CA19-9neg and CA19-9pos FC1199 at 0 hr (left) and 72 hrs (right). Scale bar = 200 µm.

Data are representative of at least two independent experiments. Data are presented as mean ± SD.

Supplemental Figure 3

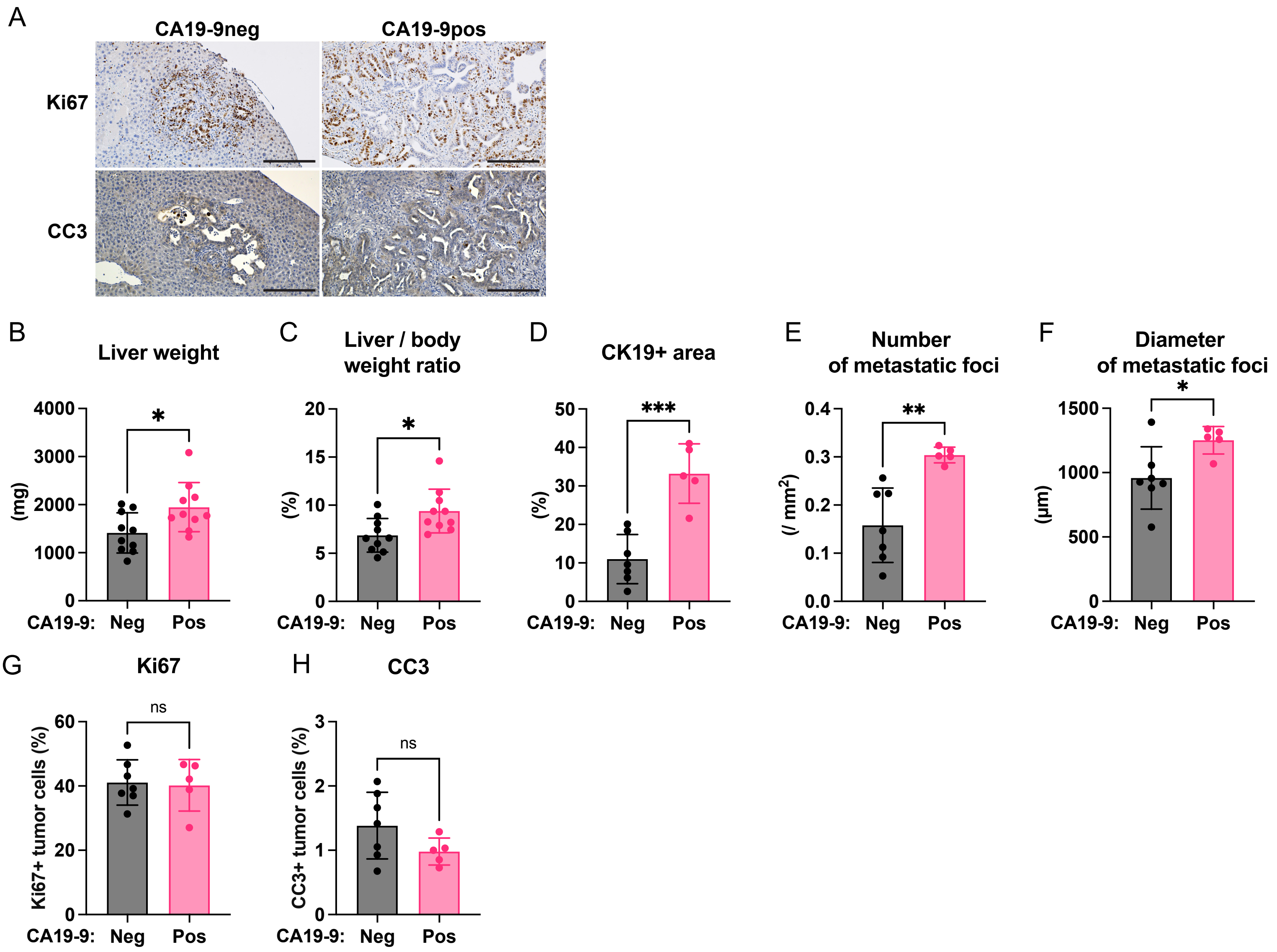


**Supplemental Figure 3. CA19-9-positive cells promote the seeding and outgrowth of the liver metastases in splenic injection models using KPCY cells.**
(A) Representative images of Ki67 (top) and CC3 (bottom) staining in liver metastases derived from FC1199 CA19-9neg and CA19-9pos cells, corresponding to the quantification shown in Figure 1I, J. Scale bar = 200 µm.
(B, C) Quantification of liver weight (B) and of liver weight normalized by body weight (C).
(D) Quantification of CK19+ area (%) across the whole liver sections.
(E) Number of metastatic foci across the whole liver sections normalized by tissue area (mm²).
(F) Mean diameter of metastatic foci measured across whole liver sections.
(G, H) Quantification of Ki67 positive cells (G) and cleaved caspase-3 (CC3) positive tumor cells (H) within liver metastasis area.
*Data are presented as mean ± SD. Each dot represents an individual mouse. Mice were injected with $1 \times 10^5$ tumor cells per mouse. Statistical significance was calculated using unpaired two-tailed t-test with Welch's correction. *P < 0.05; **P < 0.01; ***P < 0.001; ns, not significant.

Supplemental Figure 4

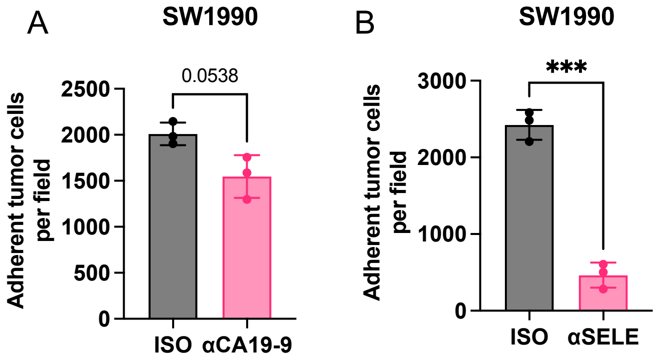


**Supplemental Figure 4. CA19-9-positive human PDAC cells exhibit E-selectin-dependent endothelial adhesion in vitro.**
(A, B) Quantification of adhered SW1990 cells treated with anti-CA19-9 antibody (5B1; αCA19-9) (A), anti-human E-selectin antibody (BBA16; αSELE) (B) compared with isotype control (ISO). Data are representative of at least two independent experiments.
*Data are presented as mean ± SD. Statistical significance was determined by unpaired two-tailed t-test with Welch's correction. ***$P < 0.001$.

Supplemental Figure 5

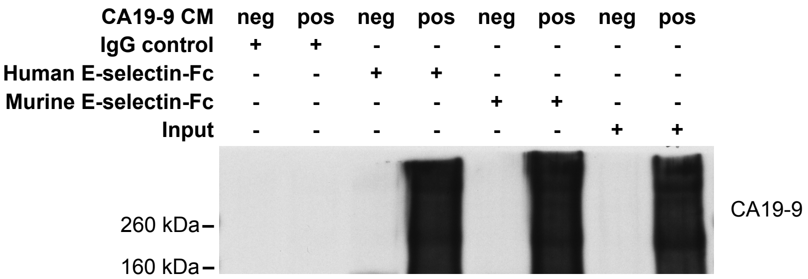


**Supplemental Figure 5. Mouse E-selectin can bind to CA19-9 in the same manner as human E-selectin.** Conditioned media were collected from CA19-9neg and CA19-9pos FC1199 cells and concentrated prior to pull-down assays. Samples were incubated with recombinant human E-selectin-Fc or recombinant murine E-selectin-Fc fusion proteins, followed by precipitation using beads. Human IgG was used as an isotype control. Immunoblotting was performed using anti-CA19-9 antibody (NS19-9).

The rightmost lanes represent input samples (concentrated conditioned media prior to pull-down). Both human and murine E-selectin-Fc demonstrated comparable binding to CA19-9 derived from CA19-9pos cells, whereas minimal signal was detected in CA19-9neg conditioned media or IgG control samples.

Supplemental Figure 6

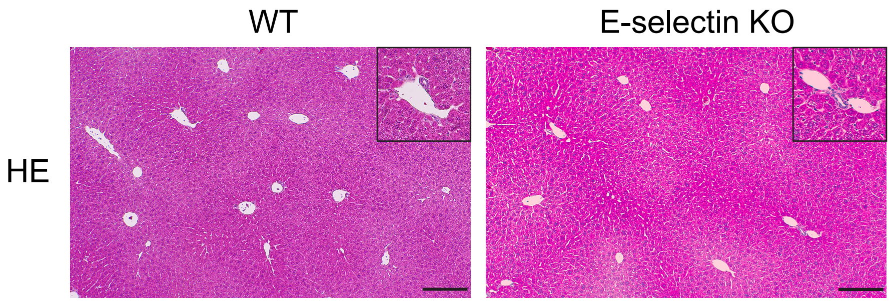


**Supplemental Figure 6. E-selectin deficiency does not induce inflammatory changes or disrupted liver architecture.**
Representative H&E images of the liver in WT and E-selectin knockout (Sele-/-) mice. Scale bar = 200 μm.

Supplemental Figure 7

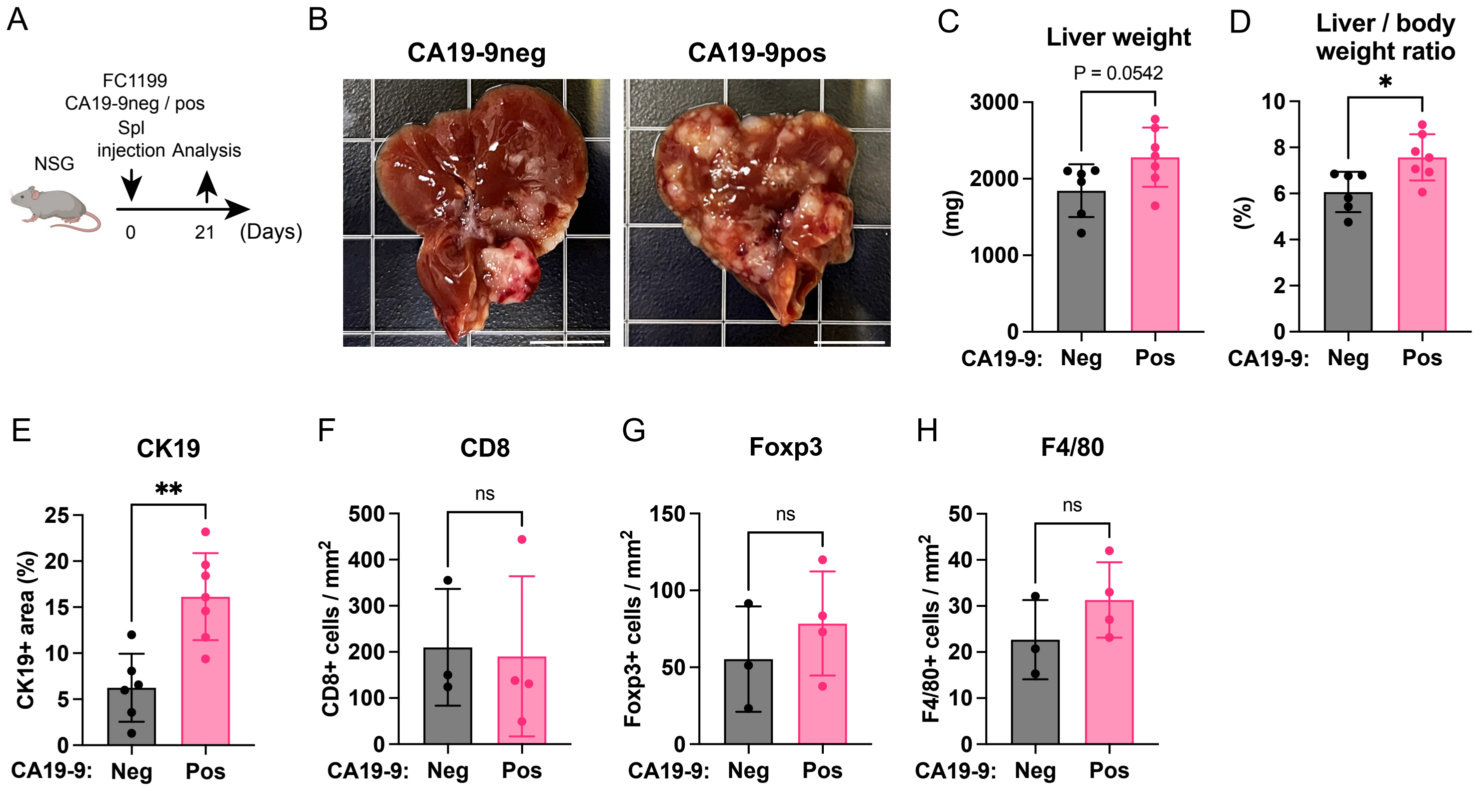


**Supplemental Figure 7. CA19-9 does not significantly alter immune cell infiltration in liver metastases.**
(A) Schema of study design in (B-E)
(B) Representative macroscopic images of the livers isolated from NSG mice injected with CA19-9neg or CA19-9pos FC1199 cells. Scale bar = 1 cm.
(C, D) Quantification of liver weight (C) and liver weight normalized by body weight (D).
(E) Quantification of CK19+ area (%) across the whole liver sections.
(F-H) Immunohistochemical quantification of CD8+ T cells (F), Foxp3+ regulatory T cells (G) and F4/80+ macrophages (H) within liver metastases in WT mice injected with CA19-9neg and CA19-9pos FC1199 cells.
*Data are presented as mean ± SD. Each dot represents an individual mouse. Mice were injected with $5 \times 10^4$ tumor cells per mouse. Statistical significance was calculated using unpaired two-tailed t-test with Welch's correction. *P < 0.05; **P < 0.01; ns, not significant.

Supplemental Figure 8

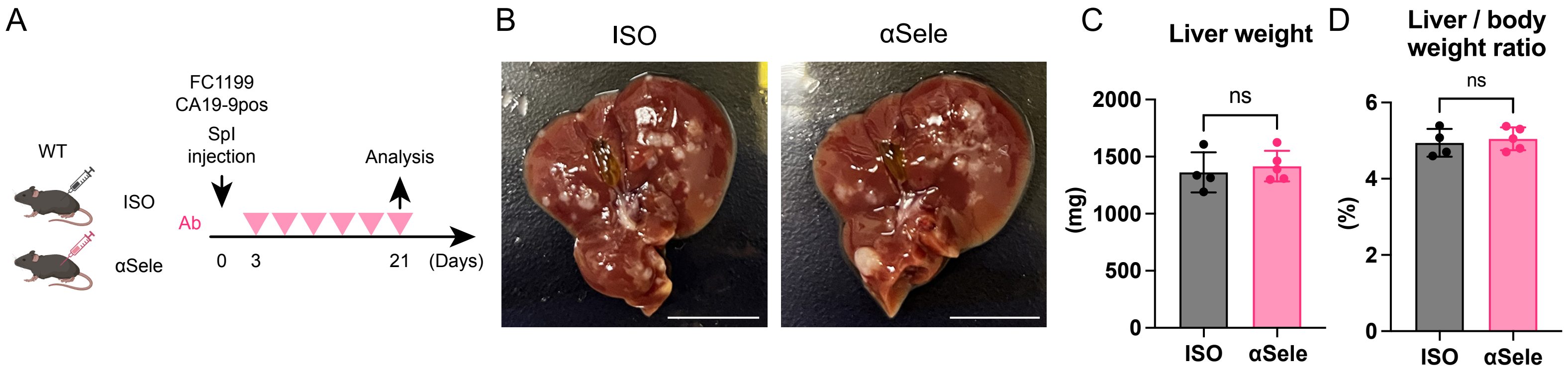


**Supplemental Figure 8. E-selectin Ab does not suppress the outgrowth of CA19-9pos liver metastasis.**
(A) Schema of study design in (B-D)
(B) Representative macroscopic images of the livers isolated from mice injected with CA19-9pos cells and treated with either isotype control (ISO) (left) or anti-mouse E-selectin antibody (9A9; αSele) (right). Scale bar = 1 cm.
(C, D) Quantification of liver weight (C) and liver weight normalized by body weight (D).
*Data are presented as mean ± SD. Each dot represents an individual mouse. Mice were injected with $1 \times 10^5$ tumor cells per mouse. Statistical significance was calculated using unpaired two-tailed t-test with Welch's correction. ns, not significant.

Supplemental Figure 9

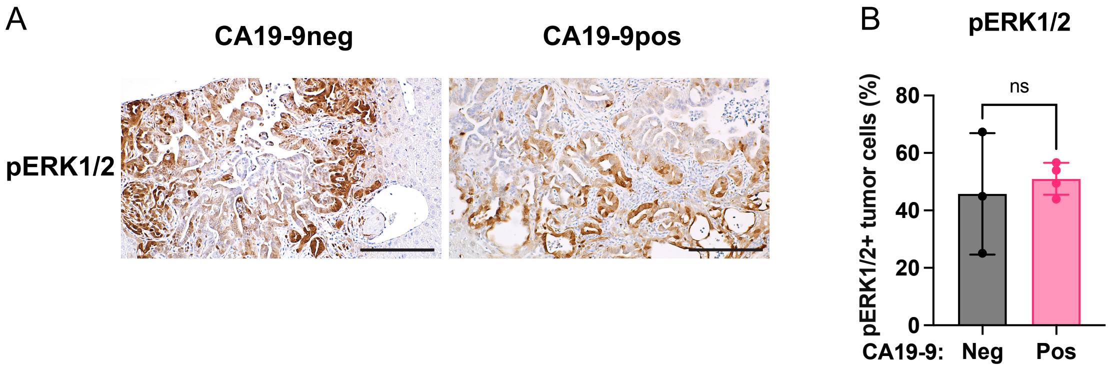


**Supplemental Figure 9. Phosphorylation of ERK remained unchanged between CA19-9neg and CA19-9pos liver metastasis.**
(A) Representative pERK1/2 IHC within liver metastases area derived from CA19-9neg or CA19-9pos FC1199 cells. Scale bar = 200 µm.
(B) Quantification of pERK1/2 positive tumor cells in liver metastasis.
*Data are presented as mean ± SD. Each dot represents an individual mouse. Statistical significance was calculated using unpaired two-tailed t-test with Welch's correction. ns, not significant.

Supplementary Table S1: Primary antibodies for IHC, IF, WB and flow cytometry

| Antibody | Clone | Company | Catalog | Application | Dilution |
|---|---|---|---|---|---|
| CA19-9 | 5B1 | MabVax | MVT-5873 | IHC / WB | 100 ng / ml |
| CA19-9 | NS19-9 | Origene | CF190083 | IF / WB / Flow | 1:2000 / 1:1000 / 1:50 |
| CK19 | TROMA-III | DSHB | TROMA-III | IHC | 1:200 |
| Ki67 | | Abcam | ab15580 | IHC | 1:2000 |
| CC3 (Asp175) | | Cell signaling Technology | 9661 | IHC | 1:200 |
| GFP (YFP) | | Abcam | ab6673 | IHC | 1:500 |
| pAkt (Ser473) | D9E | Cell signaling Technology | 4060 | IHC / WB | 1:200 / 1:2000 |
| Akt | 11E7 | Cell signaling Technology | 4685 | WB | 1:3000 |
| pErk1/2 (Thr202/Tyr204) | D13.14.4E | Cell signaling Technology | 4370 | IHC / WB | 1:200 / 1:2000 |
| Erk1/2 | 137F5 | Cell signaling Technology | 4695 | WB | 1:3000 |
| pS6 (Ser235/236) | D57.2.2E | Cell signaling Technology | 4858 | WB | 1:2000 |
| S6 | 54D2 | Cell signaling Technology | 2317 | WB | 1:3000 |
| Cofilin | D3F9 | Cell signaling Technology | 5175 | WB | 1:10000 |
| CD8 | EPR21769 | Abcam | ab217344 | IHC | 1:500 |
| Foxp3 | D6O8R | Cell signaling Technology | 12653 | IHC | 1:1000 |
| F4.80 | D2S9R | Cell signaling Technology | 70076 | IHC | 1:300 |
