## Supplemental Figure legends for "CA19-9 promotes liver metastasis of pancreatic cancer through E-selectin mediated extravasation"

**Supplemental Figure 1. CA19-9 expression in mouse and human PDAC cell lines.**

(A) Immunoblot analysis of CA19-9 expression in CA19-9^neg^ and CA19-9^pos^ murine PDAC cells (FC1199, FC1245, and KPCY).

(B) Immunoblot analysis of CA19-9 expression in human CA19-9^pos^ cell lines (hM19a 2D and SW1990). CA19-9 was detected using the anti-CA19-9 antibody 5B1.

(C) Immunoblot analysis of CA19-9 expression in Capan-2 cells with control sgRNA (sgCtrl) or FUT3-targeting sgRNA (sgFUT3). CA19-9 was detected using the anti-CA19-9 antibody 5B1.

**Supplemental Figure 2. CA19-9 expression does not alter proliferation but enhances migratory capacity *in vitro*.**

(A, B) Cell viability assay of CA19-9^neg^ and CA19-9^pos^ FC1199 (A) and KPCY (B) cells.

(C, D) Scratch wound healing assay of CA19-9^neg^ and CA19-9^pos^ FC1199 (C) and KPCY (D) cells. Wound closure was normalized to the wound area at time 0 and relative wound density was quantified.

(E) Representative images of Scratch wound healing assay using CA19-9^neg^ and CA19-9^pos^ FC1199 at 0hr (left) and 72 hrs (right). Scale bar = 200 μm.

Data are representative of at least two independent experiments. Data are presented as mean ± SD.

The rightmost lanes represent input samples (concentrated conditioned media prior to pull-down). Both human and murine E-selectin-Fc demonstrated comparable binding to CA19-9 derived from CA19-9^pos^ cells, whereas minimal signal was detected in CA19-9^neg^ conditioned media or IgG control samples.
